# Homology between the flagellar export apparatus and ATP synthetase: evidence from synteny predating the Last Universal Common Ancestor

**DOI:** 10.1101/2021.01.01.425057

**Authors:** Nicholas J. Matzke, Angela Lin, Micaella Stone, Matthew A. B. Baker

## Abstract

Evidence of homology between proteins in the ATP synthetase and the bacterial flagellar motor (BFM) has been accumulating since the 1980s. Specifically, the BFM’s Type 3 Secretion System (T3SS) export apparatus FliH, FliI, and FliJ are considered homologous to F_O_-b + F_1_-δ, F_1_-α/β, and F_1_-γ, and have similar structure and interactions. We review the discoveries that advanced the homology hypothesis and then conduct a further test by examining gene order in the two systems and their relatives. Conservation of gene order, or synteny, is often observed between closely related prokaryote species, but usually degrades with phylogenetic distance. As a result, observed conservation of synteny over vast phylogenetic distances can be evidence of shared ancestral coexpression, interaction, and function. We constructed a gene order dataset by examining the order of *fliH*, *fliI*, and *fliJ* genes across the phylogenetic breadth of flagellar and nonflagellar T3SS. We compared this to published surveys of gene order in the F_1_F_O_-ATP synthetase, its N-ATPase relatives, and the bacterial/archaeal V- and A-type ATPases. Strikingly, the *fliHIJ* gene order was deeply conserved, with the few exceptions appearing derived, and exactly matching the widely conserved F-ATPase gene order *atpFHAG*, coding for subunits *b*-*δ*-*α*-*γ*. The V/A-type ATPases have a similar conserved gene order shared for homologous components. Our results further strengthen the argument for homology between these systems, and suggest a rare case of synteny conserved over billions of years, dating back to well before the Last Universal Common Ancestor (LUCA).

## Gene order and homology hypotheses

Synteny is often observed between close relatives but tends to decay rapidly with phylogenetic distance (Lathe et al. 2000; Gómez et al. 2004; Zhulin 2017). While widely conserved operons are rare, when found, they may indicate conserved functional relationships. Co-expression, stoichiometry, and order of expression may provide preliminary evidence for the assembly and function of cooperating protein products (Dandekar et al. 1998; Gómez et al. 2004).

The observation of synteny can also be useful for proposing, or strengthening, hypotheses of homology between proteins or protein complexes. It is well-known that DNA sequence similarity between a pair of genes can eventually decay to statistical undetectability (saturation; Philippe et al. 2011); amino acid sequences, although more conserved, are subject to the same problem. When amino acid sequence similarity has decayed into the “twilight zone” (Doolittle 1986; Rost 1999), homology arguments can still be made based on conserved protein structure, function, or protein-protein interactions, but confidence will be reduced unless several pieces of evidence line up in support of the hypothesis. The discovery of synteny can support other lines of evidence in favor of a hypothesized homology and can have practical benefits as well. For example, if synteny is strongly conserved between a set of genes, this might assist identification and naming of ORFs that have been missed by automated BLAST (Lathe et al. 2000).

Furthermore, confirmation of homology between distantly-related protein complexes can inspire productive research avenues. This occurs because experimental information gathered about a well-studied protein complex – protein structure, protein interactions, functional mechanisms, etc. – provides starting points for experimental research into a less-studied, but evolutionarily related, complex. A process of “reciprocal illumination” (Hennig 1966) can ensue, where experimental work on either system helps inform work on the other, as similarities and differences are elucidated.

Here, we review a remarkable case where a sequence of homology discoveries has reciprocally illuminated research on the structure and function of two molecular machines: the F_1_F_O_-ATP synthetase (or F-ATPase) and the flagellar protein export apparatus (and their respective relatives). We add to the evidence for homology by describing shared gene order between the two systems, suggesting a case of synteny that pre-dates the Last Universal Common Ancestor (LUCA).

## The rotary motors and their relatives

For simplicity, we refer to “the” flagellum and F_1_F_O_-ATP synthetase, as they are exemplar systems that are well-studied, particularly in model systems, and because our argument can be made with reference to these. However, we acknowledge the major related systems, which we summarize below (Fig. 1).

**Figure 1.**
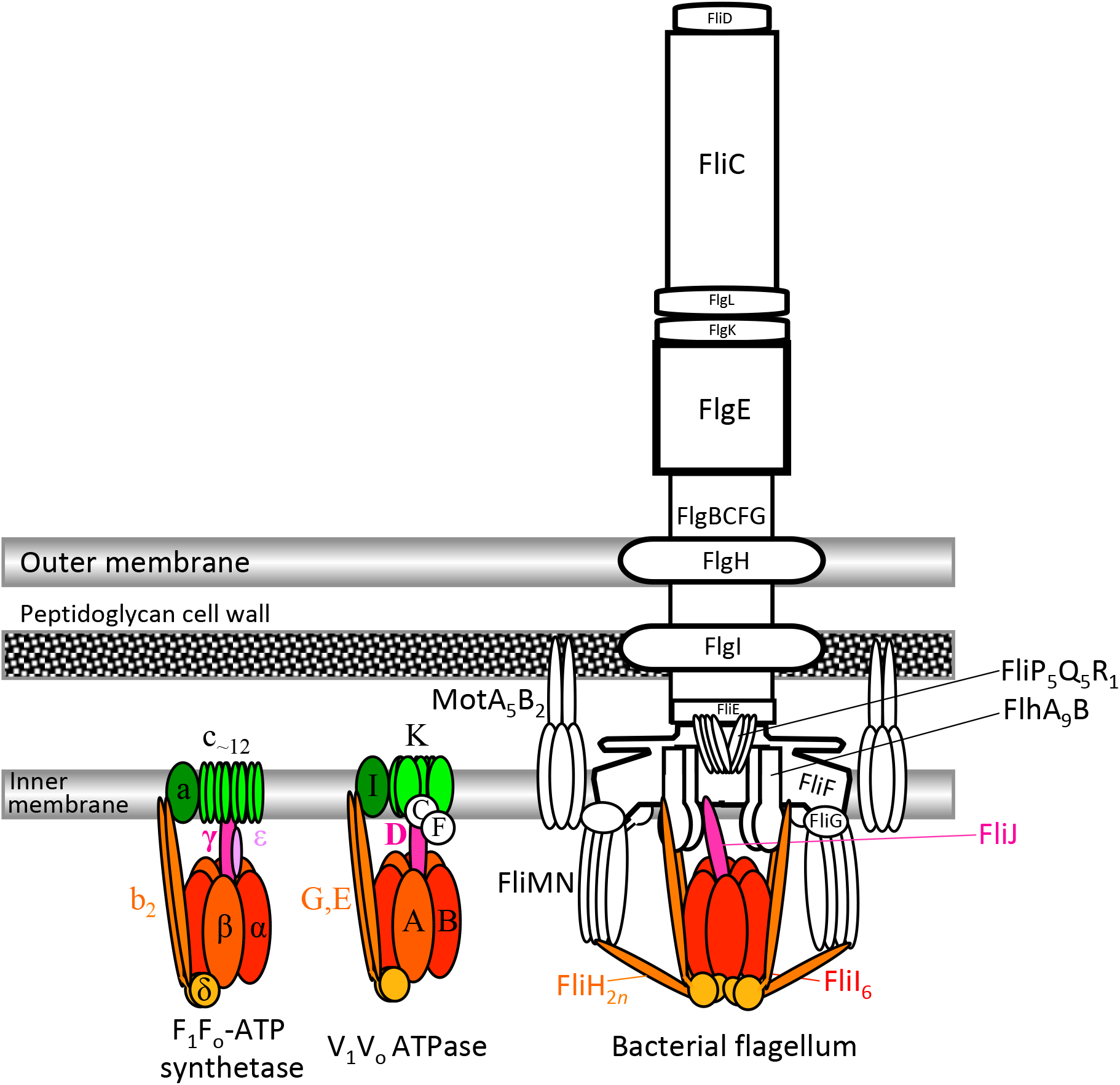
Cartoon depicting location of core structural components of F_1_F_o_-ATP synthetase, V/A-type ATPases, and the bacterial flagellum. Colored components match the colors in Fig. 2. For the V-type ATPases, naming follows the bacterial V-ATPases (Kolkema et al. 2003); see Supp. Material for corresponding names across systems. Stoichiometry is indicated with subscripts where important in this discussion, but most flagellar proteins form large oligomers (rings or filaments) with many subunits. Cartoon after Macnab (2003), updated with Minamino & Imada (2015), Minamino et al. (2017), Beeby (2020). FliH interacts with both FlhA and FliN so both positions are depicted, after Minamino & Imada (2015).

While there is no “the” flagellum, and considerable structural diversity exists across bacterial flagella (Snyder et al. 2009; Beeby et al. 2020), about 40 proteins are well-conserved across most flagella (Fig. 1; Pallen and Matzke 2006). About 11 of the proteins form a flagellar Type 3 Secretion System (F-T3SS) that exports proteins through a central channel (Milne-Davies et al 2020). The exported proteins grow a rod, hook, and flagellar filament atop the basal body. The whole structure is surrounded by an outer ring of stator/motor protein complexes (MotA_5_B_2_; Deme et al. 2020; Santiveri et al. 2020) which use the flow of H^+^ (or other cations; Ishida et al. 2019; Lai et al. 2020) to rotate the basal body (reviewed in detail by Lai et al. 2020).

The flagellar T3SS (F-T3SS) is homologous to the nonflagellar T3SSs (NF-T3SS) which form needle-like structures that secrete effector proteins into eukaryote cells; NF-T3SS are common in plant and animal pathogens. The most recent major analysis suggests that NF-T3SS are phylogenetically derived from the flagellum (Abby and Rocha 2012; Denise et al. 2020), although it appears to be a deep branch that did not originate within any particular extant bacterial phylum.

The bacterial F_1_F_O_-ATP synthetase (von Ballmoos et al. 2009; Fillingame and Steed 2014; Sobti et al. 2019; Sobti et al. 2020; Sobti et al. 2020), in contrast, is much smaller (Fig. 1). The canonical system of *E. coli* consists of 8 well-conserved proteins all localized to the inner membrane. Subunits of F_O_-c form a rotating ring in the inner membrane, and F_O_-a acts as a stator; F_O_-a and F_O_-c together form a channel allowing the flow of H^+^. The soluble F_1_-subunit consists of a heterohexameric ATPase with alternating α (noncatalytic) and β (catalytic) subunits. These are attached to the F_O_ complex with an external stalk formed by a dimer of F_O_-b, and an internal stalk formed by F_1_-γ and F_1_-∊ (refs). The stalks allow ATP hydrolysis to exert force on the F_O_-c-ring, driving rotation with respect to F_O_-a and causing proton export, or the reverse reaction (proton flow driving ATP synthesis).

Two subcategories of bacterial F_1_F_O_-ATPases are the mitochondrial (He et al. 2018) and chloroplast (Hahn et al. 2018) versions (MF_1_F_O_ and CF_1_F_O_, respectively). They each have their own experimental literature, varying nomenclature, and additional accessory subunits (Chaban 2005; Supp. Mat.), but are of relatively recent origin, being related to the F-ATPases of Rickettsiales and cyanobacteria, respectively (although many of the genes for these systems have relocated to the nuclear genome; Stoebe and Kowallik 1999; Johnston and Williams 2016). However, another subcategory, the N-ATPases, which pump Na^+^ instead of H^+^, are potentially sister to the rest of the F-ATPase clade. More broadly, the F- and N-ATPases are homologous to the archaeal A-ATPases and the V-ATPases of eukaryotic vacuoles (however, V-ATPases have since been discovered in bacterial as well, sometimes called V/A-ATPases or bacterial V-ATPases; ref). These other ATPases carry homologs to most or all of the F-ATPase subunits, although homologies are sometimes difficult to confirm due to lack of sequence similarity (Mulkidjanian et al. 2007); they also contain a variety of less-conserved accessory proteins.

## Flagellum-ATP synthetase homology: a history of discovery

The bacterial flagellum and the F_1_F_O_-ATPase are both famous for using protonmotive force to power rotary motors to do useful work. These generic similarities inspired some early scenarios that attempted to derive a flagellum directly from a synthetase (Goodenough 1998; Rizzotti 2000), but any detailed comparison of the systems reveals these as vague and fanciful (Matzke 2003).

### Homology of the ATPase proteins

The strong case for homology between the flagellum and ATP synthetase does not involve ion-powered rotary motion, but instead concerns the ATPase proteins of both systems. Early protein sequence comparisons quickly recognized highly significant amino acid sequence similarity between FliI of the flagellum, and the F_1_-α and F_1_-β subunits of the F_1_F_O_-ATP synthetase (Gogarten et al. 1989; Iwabe et al. 1989; Gogarten et al. 1992). Early discussions focused on the fact that the F_1_-ATPase’s heterohexameric structure of alternating catalytic (β) and non-catalytic (α) subunits proteins is also found in the V- and A-ATPases, suggesting that the heterohexameric ATPase was found in the Last Universal Common Ancestor (LUCA). The catalytic and non-catalytic subunit were derived by gene duplication from a common ancestor that was a catalytic homohexamer (as in more distant relatives like the transcription termination factor Rho, and the P-loop ATPase RecA; Mulkidjanian et al. 2007). Initial interest focused on the suggestion that these ancient paralogs could thus be used to provide outgroups for each other, allowing rooting of the phylogenetic Tree of Life (ToL) between the bacteria and the archaea/eukaryote clade (Gogarten and Kibak 1992; Shih and Matzke 2013; Shih et al. 2017). While conceptually appealing, later work has questioned the idea the small collection of pre-LUCA paralogs retains sufficient signal to estimate the ToL root or the premise that different genes would record the same organismal history in the first place (Matzke et al. 2014; Gouy et al. 2015).

### Homology of the outer stalk

Surprisingly, although the homology between FliI and the F_1_-α/β was recognized early, it was not until Jackson & Plano (2000) that it was suggested some flagellar proteins might form a multiprotein complex similar in structure to F1 and V1, inspired by BLAST hits between the FliH homolog YscL and the external stalk protein V1-e (Ge et al. 1996 also noticed sequence similarity between *Borrelia* FliH and F_O_-b). Only in 2003 was homology deployed to inform models of flagellum structure. Prior to this date, cartoons of flagellum structure represent FliI as a monomer proximal to the inner membrane. However, in Blocker et al. (2003) hypothesized, on the basis of F_1_-ATPase homology, that the *Shigella* NF-T3SS Spa47 (a FliI homolog) formed a homohexamer. Claret *et al.* (2003) showed hexamerization of FliI with electron microscopy.

Further observations of FliH/F_O_-b sequence similarity, and additional similarities (both FliH and F0-b dimerize, both associate with ATPases (FliI, F_1_-α/β), and both associate with membrane proteins via N-terminal α-helices) inspired suggestions that a FliH dimer might serve an external stalk role similar to F_O_-b (Matzke 2003; Pallen et al. 2005; Lane et al. 2006). Lane et al. (2006) also found analogies between F_1_-δ and the globular, C-terminal domain of FliH; F_1_-δ forms the membrane-distal part of the F_1_F_O_-ATPase external stalk, and both FliH-C and F_1_F_O_-δ bind to amphipathic helices at the N-terminus of FliI and F_1_-β, respectively. They summed up the similar interactions between FliHI and F_O_-b, F_1_-δ, and F_1_-α/β as having “uncanny similarity,” although they termed them analogous, rather than homologous.

Pallen et al. (2006) used PSI-BLAST to conduct a systematic search for distant homologs of FliH/YscL and found remote, but significant, sequence similarity (~18-20%) to F_O_-b and the external stalk E subunit of V-ATPase. These matches occurred with just the N-terminal ~115 residues of FliH/YscL, so Pallen et al. conducted a separate search with the C-terminal region (YscL positions 115-223). Variations on these searches yielded repeated, although nonsignificant (expected number of hits E-value ranging from 0.14-8.5) hits to the F_1_-δ subunit (known as the OSCP in the MF_1_Fo-ATPase). They buttressed the argument by presenting highly suggestive multiple sequence alignments, and by noting that the genes for F_O_-b and F_1_-δ (atpF and atpH in *E. coli*) are often adjacent, and that natural fusions of F_O_-b and F_1_-δ are observed in some bacterial genomes.

The homology of three F-ATPase proteins to two flagellar export proteins inspired exploration of possible implications, leading Pallen & Matzke (2006) to note “the existence of ancient structural and functional similarities between these two types of molecular rotary motor, which we predict will become ever more apparent as work on these systems progresses.”

Mulkidjanian, et al. (2007) reviewed the similarities among V/F/A-ATPases, flagellar and nonflagellar T3SS, but found no homology between the central stalks proteins (FliJ/YscO in T3SS; F_1_-γ in F-ATPases, and V_1_-D/F in V-type ATPases). However, they note the membership of FliI/F_1_-α/β in the vast group of P-loop NTPases (Lupas and Martin 2002), specifically within the RecA-like motor ATPases (Ye et al. 2004), and even more specifically their relationship to Rho, the transcription-termination factor that forms a homohexamer ring to act as an ATP-powered RNA helicase. They propose that a cytoplasmic RNA helicase could associate with a transmembrane channel (the proto-F_O_-c ring) to become an RNA translocase, and later transition into a protein translocase with the addition of an external stalk. From here, they propose the divergence of the VFA-ATPases and the T3SS. On the VFA-ATPase line, ion translocation (Na+ or H+; Mulkidjanian 2009 argue for Na+ as the easier starting point) is added for unclear reasons. Then, the central stalks (F_1_-γ, V1-D) evolved independently from translocating proteins that became temporarily trapped and instead rotated the proto-c-ring, driving ion translocation in reverse, coupling ATP synthesis/hydrolysis with an ion gradient. Permanent recruitment of these proto-stalk proteins and specialisation on the ion-pumping/ATP synthase function results in the modern systems. The further evolution of the proto-T3SS is not detailed, although it is suggested that FliI still acts directly in the protein translocase activity, threading polypeptides through the middle of the hexamer.

### Homology of the inner stalk

Additional speculation about the descent of T3SS and V/F/A-ATPases from a common ancestral system accumulated (Imada et al. 2007; Mulkidjanian et al. 2009) until the next major advance homologizing the systems. This was the crystallization of FliJ (Ibuki et al. 2009), leading to a solved 3D structure that was immediately recognized as highly similar to F_1_-γ and V1-D (Ibuki et al. 2011). Both proteins consist of two long antiparallel α-helices. In spite of no significant sequence similarity, structure-guided sequence alignment indicated conserved residues in regions involved in interaction with other proteins. Strikingly, cryo-EM images showed FliJ localizing off-center within the FliI hexamer ring; in other words, “[t]he FliI6FliJ complex looks just like F_1_-ATPase” (Ibuki et al. 2011). Similar results were soon found for the NF-T3SS homolog of FliJ, EscO/YscO (Romo-Castillo et al. 2014). Additional similarities accumulated. Ibuki et al. (2011) noticed a conserved binding site between FliJ and F_1_-γ corresponding to where F_1_-γ interacts with F_1_-∊. Ibuki et al. (2013) showed that the corresponding residues of FliJ are involved in interactions with FlhA-L, a possibly α-helical linker region between the N-terminal transmembrane helices of FlhA and the cytoplasmic C-terminal domain. A FlhA nonamer ring forms the core of the membrane-bound portion of the T3SS, suggesting at least a general similarity to the F_O_ subunit, and demonstration that FliJ stretched from inside the FliI ring to a membrane-bound protein finally clinched the case that the FliI hexamer took the same orientation with respect to the inner membrane as F_1_-α3β3 (that is, FliI C-terminal region faces the membrane, and N-terminal faces away; (Ibuki et al. 2011).

Finally, Kishikawa et al. (2013) tested whether FliJ could rotate like F_1_-γ, constructing a chimeric complex of *Salmonella* FliJ with *Thermus aquaticus* V1-A3B3, finding that FliJ localized to the center of the ring and rotated bidirectionally. Diepold & Armitage (2015) suggested that “the FliJ/SctO stalk might rotate,” and incredibly, a chimera of FliJ with the 21 C-terminal residues of V1-D was shown to rotate unidirectionally inside the V1A3B3-ATPase in the presence of ATP at a speed comparable to V1-D (Baba et al. 2016). Rotation, though weaker, was also shown for a F_1_-FliJ chimera (Baba et al. 2016), suggesting torque generation through a coarse-grained, non-sequence-specific interaction that must date back before the LCA (Noji et al. 2017).

The discoveries mentioned above indicate that the homology hypothesis has served as a productive research avenue (as suggested by e.g. Pallen et al. 2006; Pallen and Matzke 2006), and homology between the F_1_-ATPase and FliHIJ is now well-accepted and routinely considered in studies of both structural/functional studies of both systems. However, it should be kept in mind that, apart from the highly significant sequence similarity between FliI and F_1_-α/β (and many more distant relatives; Iyer et al. 2004; Ye et al. 2004), which are large proteins with conserved sites of catalysis and conserved structure at all levels, the other proteins are small and structurally simple, formed primarily of extended α-helices. The sequence similarity evidence, judged by e-values, is weak (FliH/F_O_-b+F_1_-δ, particularly the latter) or non-existent (FliJ/F_1_-γ). And, while the case for homology is reasonably strong *in toto*, as a large number of similarities, individually only suggestive, point in the same direction, it is somewhat conceivable that they might be explained away by convergence. Furthermore, experimental work suggests the possibility of some substantial dissimilarities. For example, while experiments and homology have suggested that FliH_2_ dimerizes to form the outer stalk in a fashion similar to F_O_-b, cryoelectron tomography of the NF-T3SS of *Shigella flexneri* indicates that the FliH homolog (MxiN) forms six “spokes” that link the N-terminal regions of the FliI hexamer subunits to FliN subunits, of the C-ring, capping the T3SS structure (Hu et al. 2015; Gao et al. 2018). The functions of F_1_-γ and FliJ also appear quite different; F_1_-γ rotates with the F_O_-c ring, coupling ATP synthetase activity to ion flow, whereas FliJ seems to play a role in making substrate proteins (axial proteins, including flagellin and rod/hook/linker/cap proteins and NF-T3SS equivalents) export-competent by removing chaperones (chaperones FlgN, FliS, FliT are all structurally similar; Khanra et al. 2016) and delivering substrate to the ion/protein antiporter via interactions with FlhA (Gao et al. 2018; Minamino et al. 2020).

### Another source of evidence: Gene order

We investigated whether gene order evidence might strengthen or weaken the hypothesis of homology between FliHIJ and F_1_-ATPase. Gene order has occasionally been used to identify homologous proteins in T3SS. For example, FliJ homologs in NF-T3SS (Romo-Castillo et al. 2014) as the gene for the FliJ homolog is typically downstream from the gene for the FliI homolog (Evans et al. 2006). Similarly, Pallen et al. (2006) noted that the genes for F_O_-b and F_1_-δ are typically adjacent (Wilkens and Capaldi 1998) and sometimes fused (Féthière et al. 2004; Pallen et al. 2006). While *fliH*, *fliI*, and *fliJ* were named alphabetically in order in the unified flagellar gene nomenclature established by Iino et al. (1988), this scheme was based on *E. coli* & *Salmonella* sp. Liu & Ochman (2007) surveyed operon structure and gene order across many flagellar systems and found that the alphabetical ordering broke down at larger phylogenetic scales. They did note that certain small clusters were usually adjacent, e.g. *flgBC*, *flgKL*, *flhBA*, *fliMN*, *fliPQR*, but they did not mention *fliHIJ*. This was probably because the *fliHIJ* cluster was missing from 4 of the 10 groups they summarized in their Figure 3. The *fliJ* gene was thought to encode one of several flagellar chaperons, and Liu and Ochman stated that this “auxiliary” gene had a “sporadic” distribution in their dataset. This was likely due to failure to detect FliJ homologs due to its less-conserved sequence.

The study of gene order has been more systematic in F- and V/A-type ATPases, so we are able to rely on previously-published reviews for these systems. Gene order in the F_1_F_O_-ATPase and relatives is highly, but not universally, conserved across bacterial phyla and divergent systems (Koumandou and Kossida 2014; Niu et al. 2017). The same is true for V/A-type ATPases in bacteria and archaea (Lolkema et al. 2003).

## Methods

We surveyed the order of the *fliH*, *fliI*, and *fliJ* genes across bacteria. We chose species representing each major flagellated clade/phylum (Fig. 2), using Snyder et al. (2009) for guidance. Our strategy was to identify *fliI* homologs, as FliI is well-conserved and usually identified by automated annotations, and then identify the flanking genes. For each species, we searched Ensembl’s bacterial genome browser (Howe et al. 2020). We started by searching by name for FliI/YscN. If that failed, we searched for FliF, which is also highly conserved and sometimes annotated when FliI is not, and then searched neighboring genes for a FliI homolog. If both of those approaches failed, we BLASTed the genome with the *E. coli* FliI amino acid sequence.

**Figure 2.**
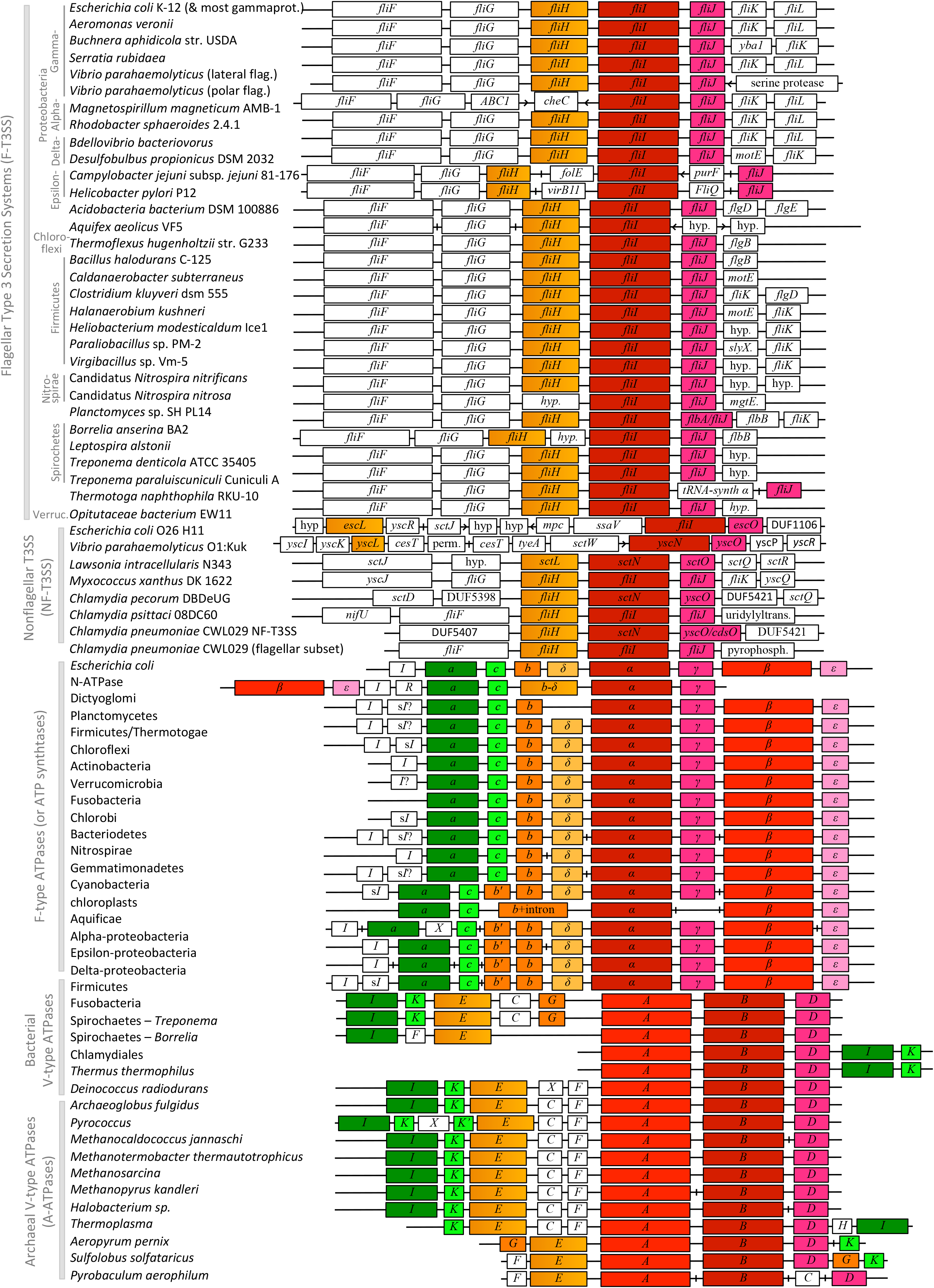
Conserved gene order between *fliHIJ* genes and V/F/A-ATPase genes. Homologs of *fliH*, *fliI*, and *fliJ* are colored orange, red, and dark pink, respectively. Homologs of V/F/A-ATPase a-, c-, and ε-subunits are dark green, light green, and light pink. Other genes are white. Genes are ordered from upstream to downstream with respect to the *fliI* exon. Genes coded on the opposite strand are indicated by left arrows (<); return to the *fliI* strand by right arrows (>). Large breaks are indicated by vertical lines (|) as in Lolkema et al. (2003). For T3SS gene order, see Methods and Supplemental Material; Supp. Mat. also includes the full names of strains. F-ATPase genes are named by the protein subunit, with the order taken from Koumandou & Kossida (2014). V/A-ATPase gene names & order from Kolkema et al. (2003). *E. coli* F_1_F_O_-ATPase after Nielsen et al. (1984). Chloroplast (*Arabidopsis*) gene order from https://www.ncbi.nlm.nih.gov/gene/844790; chloroplast *atpG* (b’, II), *atpC* (γ), and *atpD* (δ), are encoded in the plant nuclear genome (Hermans et al. 1988; Stoebe & Kowallik 1999). This is a schematic depiction to highlight gene order; the gene lengths and spaces between genes are not intended to be exact. *Notes:* N-ATPase “b” is a b+δ fusion (Dibrova et al. 2010), as is V1-E (Pallen et al. 2006). The V_O_ I-subunit is homologous to F_O_–a and the subunit named “a” in yeast V-ATPases (Chaban et al. 2005). The F_O_–I subunit (and the homologous sI) are Ca^2+^/Mg^2+^ transporters, unique to F-ATPases (Koumandou & Kossida 2014). DUF: conserved Domain of Unknown Function; hyp. = hypothetical protein in database, not identified as homologous to T3SS genes by the database or manual searches.

Once *fliI* was located, we identified the flanking genes using the following approach. If the protein was automatically identified, we used that label. If it was not, we checked the protein family and then conserved domain information, using the first identification found. If these approaches revealed nothing, we downloaded the amino acid sequence and used PSI-BLAST (Altschul et al. 1997) on the RefSeq Select proteins database (refseq_select; Li et al. 2020). If all of those approaches failed, we recorded the protein as a hypothetical protein, or a DUF (conserved Domain of Unknown Function), if so identified in the database. To provide genomic context, we used the same approach on the 2 genes upstream and downstream of the *fliHIJ* cluster, recording the flanking 2 genes on each side of *fliHIJ* if they were flagellar, or 1 if the further genes were nonflagellar.

Gene identifications were recorded in transcription order from upstream to downstream, ordered with respect to the *fliI* homolog. Cases where coding switched to the opposite strand from *fliI* were indicated with “<”. In cases where *fliH* or *fliJ* was not found near *fliI*, we BLASTed the genome with the FliH or FliJ amino acid sequence from a relative. Once the homolog was identified, we identified the flanking 2 genes on each side if they were flagellar, or 1 if the further genes were nonflagellar. The non-neighbor *fliH/J* genes are displayed in Fig. 2 in the same order as the canonical *fliHIJ* ordering, but with “|” indicating a discontinuity meaning that the genes are not in the same neighborhood.

F-ATPase and prokaryote V/A-ATPase gene orders were taken from Fig. 4 of Koumandou & Kossida (2014) and Table 2 of Lolkema et al. (2003), using their clade names and gene names. The Koumandou & Kossida (2014) dataset represents summaries of the most common gene arrangements in each clade according to their analyses; Lolkema et al. (2003) uses representative species. We did not investigate gene order in eukaryotic V-ATPases as eukaryotic genome organization is vastly more complex and presumably derived with respect to the A-type systems.

## Results

Figure 2 depicts gene order across highly divergent representatives of F-T3SS, NF-T3SS, F-type ATPases, and V/A-type ATPases from bacteria and archaea. A high degree of conservation of gene order of the *fliHIJ* homologs can be observed across the T3SS without sophisticated analysis. While gene order breaks down for some Alpha- and Epsilon-proteobacterial systems, as noted by Liu & Ochman (2007), it is otherwise conserved across proteobacteria as well as deep splits in the bacterial tree, including Chloroflexi, Spirochetes, Firmicutes, Nitrospirae, Thermotogae, and Verrucomicrobia. The gene order is also found in the presumed “flagellar remnant” (Betts-Hampikian and Fields 2010) system found in Chlamydiae in addition to their classic NF-T3SS; some marine Chlamydiae are now known to have flagella (Collingro et al. 2017; Dharamshi et al. 2020).

While the conserved gene order is disrupted in *E. coli* and *Vibrio* NF-T3SS by the relocation of *escL*/*yscL* (*fliH* homologs), the standard gene order is retained across the deeper branches of NF-T3SS, including *Myxococcus* and *Chlamydia* lineages. Overall, whatever exact phylogeny is assumed to represent the relationships between the different bacterial phyla (a difficult question; Ishida et al. 2019) and the NF-T3SS (Abby and Rocha 2012), it is clear that *fliHIJ* represents the ancestral gene order.

Turning to the F-type ATPases, the gene order found in the model E. coli system is conserved with rare exceptions across phyla; Koumandou & Kossida (2014) conclude it is the ancestral gene order. Strikingly, the genes for F_O_-b, F_1_-δ, F_1_-α, and F_1_-γ are arranged identically with their *fliHIJ* homologs, taking into account that F_O_-b and F_1_-δ are considered to be homologous to the N-terminal and C-terminal regions of FliH, respectively.

The prokaryotic V/A-ATPases also show strong conservation of gene order, with I-K-E-C-G-A-B-D representing the likely ancestral arrangement (Lolkema et al. 2003), with the poorly-conserved F (and the associated nomenclatural nightmare; Lolkema et al. 2003) appearing in various locations. In 2003, Lolkema et al. assumed that subunit E formed the central stalk and that D was the peripheral stalk, so the similarity to the F-type ATPase gene order was not apparent.

However, it is now recognized that E and G are external stalks (Vasanthakumar and Rubinstein 2020; see Supp. Mat. for nomenclatural conversions between prokaryotic and eukaryotic systems), where E corresponds to F_O_-b and F_1_-δ, and G is a shorter external stalk corresponding to F_O_-b (Pallen et al. 2006). Subunit D is the central stalk, homologous to F_1_-γ (reviewed above). The major exceptions to the conserved gene order are the variable positions of the central stalk components F and C. The C-subunit is named “d” in the eukaryotic systems (Féthière et al. 2004; Chaban 2005) and forms the interface between the central stalk and the ion-transducing membrane ring (V_O_-K, corresponding to F_O_-c).

Examination of the order of the genes for the ATPase proteins with respect to the central stalk gene shows that the V_1_-B subunit (noncatalytic and considered homologous to F_1_-α) is almost always directly upstream of the gene for V_1_-D. This corresponds with the arrangement of F_1_-α and F_1_-γ. However, the order of the catalytic and noncatalytic subunits is reversed between the V_1_ (AB) and F_1_ (αβ, with γ in-between).

## Discussion

The gene orders of the *fliHIJ* homologs are observed to be highly, although not universally, conserved, across flagellar and nonflagellar T3SS and the prokaryotic F- and V/A-type ATPase systems. As the duplication of an ancestral FliI-like ATPase to produce the paralogous catalytic F_1_-β/V_1_-A and noncatalytic F_1_-α/V_1_-B clades predates the LUCA (Iwabe et al. 1989; Shih and Matzke 2013), and the divergence of FliI must precede that, this constitutes the most ancient case of synteny involving more than 2 genes of which we are aware.

Our results suggest a parsimonious scenario to explain the ancestral gene orders in the studied systems. The proto-F_1_-like system was encoded by an operon with genes coding for the equivalents of F_O_-b, F_1_-δ, FliI, and FliJ. Duplication of the FliI homolog produced 2 catalytic subunits in tandem. The second of these (representing proto-F_1_-α/V_1_-B) lost catalytic ability, and then after the divergence of the F- and V-type systems, the ancestor of F_1_-β was transposed to be just downstream of F_1_-γ.

In the same scenario, the external stalk subunits originated by (1) fusion of the F_O_-b and F_1_-δ subunits to produced proto-FliH in the F-T3SS ancestral lineage; (2) retention of the ancestral arrangement in the F-ATPase lineage, with occasional later duplications of F_O_-b and fusions with F_1_-δ; and (3) a similar process of duplication and fusion producing the E and G external stalk subunits in the V-ATPase lineage, along with the addition of other subunits associated with the central stalk.

Several implications follow from our results. First, conserved synteny further strengthens the case for homology between the two systems, and the specific homologized proteins. Although there was little remaining doubt in the literature concerning these homologies, additional evidence helps to confirm research is on the right track, and adds genomic context to inferences about ancestral complexes based on shared sequence, structure, and functional detail.

Secondly, our results confirm earlier conclusions, based only on operon structure, that the F-type ATPases are made up of evolutionary submodules (Niu et al. 2017); indeed, these submodules appear to sit even deeper into evolutionary history than Niu et al. suggest.

Thirdly, they constitute a dramatic illustration of the hypothesis that widely-conserved gene order in prokaryotes indicates conserved protein-protein interactions (Dandekar et al. 1998; Lathe et al. 2000). In addition, the fact that exceptions to the conserved gene order are known and thus are possible, suggests that systems with alternative arrangements have somehow compensated for what is otherwise a common, conserved mechanism for regulating gene expression and stoichiometry across these systems.

Fourthly, cases of highly-conserved gene order suggest an obvious way to identify gene products in prokaryote genomes, when ORFs are unidentified by automated approaches: if fliI is identified, then fliH and fliJ are likely the flanking genes, even if the sequence similarity in these short genes has decayed beyond detectability. For example, we expect that the hypothetical gene upstream of *fliI* in *Planctomyces* sp. SH PL14 (Fig. 2) will turn out to be a *fliH* homolog, even though PSI-BLAST searches using the amino acid sequence turned up only a conserved, but unidentified, protein family.

Finally, discoveries of homology suggest avenues for further homology searches. It is tempting to postulate that the conserved F_1_-like complex suggests an ancestral association with a membrane-bound system (Mulkidjanian et al. 2007; Mulkidjanian et al. 2009), and that therefore at least parts of the membrane-bound components of the T3SS export apparatus should be ancient homologs of the F_O_ subunits. However, despite the fact that experimental discoveries have indicated certain unexpected similarities between the systems – notably, the T3SS uses ion-motive force to power protein export – Beeby et al. (2020) report no homology with the FliPQR proteins. These proteins were long strongly predicted to be transmembrane proteins, but solved structures indicate they form a gate that is withdrawn above the inner membrane to bind with FliE, the beginning of the axial filament. Beeby et al. thus conclude that there is no homology between T3SS_s_ and F_O_, and the F_1_-like complex was a late addition to the flagellar export system. They support this by noting that the FliHIJ complex is not absolutely required for protein export, which can be driven by ion motive force alone, although ATPase activity it makes it highly efficient.

Our results indicate a few points for consideration. First, it is a mistake to assume that either the extant systems in either the T3SS or V/F/A-ATPase branch represent “the ancestral” configuration. Rather, both systems retain a mixture of ancestral and derived features of gene order. This principle will likely apply to other homologies between the systems; for instance, it is possible that the F_O_ subunit is substantially simplified and specialized from a larger and more complex protein-translocating ancestor (Mulkidjanian et al. 2007). In addition, the fact that the ancestral F_1_-like subunit contained both external and internal stalks suggests a multi-subunit interactive function that might not be easily transposed into a new system. We suggest caution when evaluating possible indications of homology between F_O_ and parts of the T3SS. It is possible that homology might appear in retained interactions (e.g., FliJ interacts in part with FlhA-L, the linker region between the N-terminal transmembrane and C-terminal cytoplasmic domains of FlhA; Ibuki et al. 2013) and molecular mechanisms (e.g., the conversion of ion-motive force into conformational changes), even if the three-dimensional structures have been highly modified.

## Acknowledgements / Funding

MABB is supported by the UNSW Scientia Research Fellowship, the CSIRO Synthetic Biology Future Science Platform 2018 Project Grant and ARC Discovery Project DP190100497. NJM is supported by the University of Auckland Faculty Research Development Fund (FRDF) Strategic Initiative - Early Career Researcher Fund, Project #3722433, and New Zealand Marsden Grants 16-UOA-277 and 18-UOA-034. MS was a summer student volunteer at the University of Auckland.

